# Modeling Fast CICI Calcium Waves

**DOI:** 10.64898/2026.02.12.705545

**Authors:** Z. Peradzyński, B. Kaźmierczak, S. Biaźecki

## Abstract

Following the suggestion of L. F. Jaffe [1] we propose a mathematical model of fast calcium induced calcium influx waves (CICI Waves). They can propagate at relatively high speeds (up to 1300 micrometers/s). According to [**1**], they propagate due to a mechanochemical interaction of actomyosin network with the cell membrane. The local stretching of the membrane caused by actin filaments opens mechanically operated ion channels resulting in the influx of calcium to the cell. Moreover, stretching a cell’s membrane at one point opens nearby stretch activated calcium channels because the mechanical force is relayed by the actin filaments interconnected by myosin bridges. The number of bridges as well as filament density increases with calcium concentration, causing the contraction of the actomyosin network. Thus, the force acting on the membrane from tangled actin filaments is transmitted ahead of the moving front of the calcium concentration. As a result, the ion channels are opened even before the signal of calcium reaches them. This leads to much larger propagation speed of CICI waves in comparison with calcium induced calcium released (CICR) waves, where the wave is sustained by the diffusion of calcium and autocatalytic release of calcium from the internal stores (e.g. endoplasmic reticula).

## 1. Introduction

Since the experimental discovery of calcium waves in 1978, observed in the Medaka egg after fertilization [2], an enormous effort was put in the elucidation of underlying mechanisms. There are several types of waves, which can propagate within the cells (intracellular waves), across the tissue (intercellular waves) or through whole organs [1,3]. In the normal state of the cell the calcium concentration is rather low, however after sufficiently strong local chemical or mechanical stimulation, a concentration of calcium can locally increase so as to trigger propagation of a wave of calcium concentration. In an idealized long biological cell (like a myocyte cell) such a wave would take a form of a pulse of calcium concentration moving with a constant speed. Among various types of waves of calcium concentration which are observed in living cells, the fast waves or calcium induced calcium release (CICR) are probably best known [**3 - 10**]. Their propagation is supported by diffusion of calcium in the cytosol and the autocatalytic release of calcium from the internal stores (e.g. endoplasmic reticulum). CICR waves are supported by the autocatalytic mechanism. If the calcium concentration in the cytosol is slightly bigger than the equilibrium concentration (ground state), then calcium is absorbed into intracellular stores (endoplasmic reticulum and mitochondria); thus the equilibrium is restored. However, when the calcium concentration in the cytosol exceeds a certain threshold value, then the calcium is released from the stores. This process may continue until the concentration reaches the upper (quasi-stable) equilibrium. This supply of calcium from the internal stores sustains the CICR waves propagation. The simplest mathematical model is based on a single reaction diffusion equation for calcium concentration *c*(*t, x*)

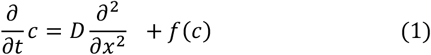

with a bistable reaction term *f*(*c*). Here, *f*(*c*) = 0 defines the possible equilibria. This equation has travelling wave solutions of heteroclinic type i.e., fronts connecting two stable equilibria. For example, for

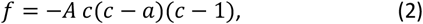

where 0 < *a* < 1 these solutions are given (up to the shift *x* → *x* + *x*_0_) by [3] :

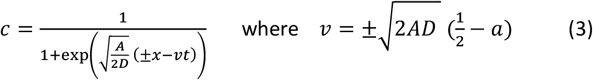

where the propagation velocity is 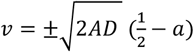. For simplicity we have assumed that the first equilibrium concentration is zero although the basic concentration in the cytosol although small is greater than zero. *c* = 1 is the other stable equilibrium, whereas *a* is the intermediate unstable equilibrium. One can easily rescale Eq.(1) to have *c*_0_ as the first equilibrium putting 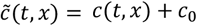 in Eqs.(1-2). Then the source term becomes 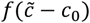 and the new equilibria are equal 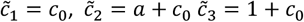. Let us notice that for *c* < *a* the source term (2) is negative; the excess of calcium is evacuated by special pumps into internal stores. However, for *c* > *a* a complex mechanism is switched resulting in an autocatalytic release of calcium until the maximal level c=1 is reached. The wave described by Eq. (3) for *t* = *const* represents a monotone front vanishing at infinity (if “ +” sign is chosen) and approaching *c* = 1 for *x* → −∞.

### Remark

Let *f*(*u*) be a bistable function and let *f*(*K*) = 0 for *K* ≠ 0 us notice that if then we introducing the new variable *U* = *u*/*K* we transform *f* to a new bistable function 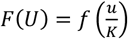, such that *F*(1) = 0. We have useful property, which can be easily proved by inspection:

### Theorem

Let *F* be a bistable function having two stable equilibria *U*_1_ and *U*_3_ = 1 (and unstable between), where *U*_1_ < 1, i.e. *F*(*U*_1_) = 0, *F*(1) = 0 and *F*_*U*_(*U*_1_), *F*_*U*_(1) < 0. Let *U*(*x*) be a heteroclinic solution of

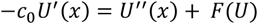

which connects these two stable equilibria of *F*, and let 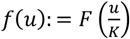, then 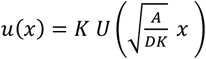 and 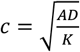 is a heteroclinic solution of −*cu*^′^(*x*) = *Du*^′′^(*x*) + *Af*(*u*) connecting stable equilibria *u*_1_ = *KU*_1_ and *u*_2_= *K* of *f*(*u*). In other words, 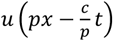 with 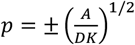 is the heteroclinic traveling solution of the following reaction diffusion equation.

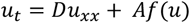

In reality however, calcium waves are homoclinic pulses with sharp front and slowly falling rear. The autocatalytic release is switched of and by action of special pumps calcium is moved back to internal stores. In this way the lowest equilibrium concentration (ground state) is restored. Therefore solutions of Eq.(1) can describe properly only the front part of a real calcium wave. The typical speed of CICR fast waves is in the range 10-30 μm/s. It appears however [**1**], that in certain cells there are waves of calcium concentration with much higher speeds, 100 -1300 μm/s. As noticed by Lionel Jaffe the CICR mechanism cannot explain so high speeds. In addition, these amazingly fast waves are observed in some cells which are not equipped with internal stores of calcium. This clearly suggest that calcium comes from the extracellular space. In [**11**] authors prove that the calcium waves activating the eggs observed in Drosophila oocytes are supported by external sources of calcium. The calcium is delivered from extracellular space through mechanosensitive ion channels located in the membrane. Following Jaffe, we will speak of calcium induced calcium influx (CICI) waves. According to L. Jaffe hypothesis, the crucial role in supplying calcium to the cell cytosol is played by stretched activated ion channels. The local stretching is caused by actin filaments anchored in the membrane.

When calcium concentration increases the network of actin filaments contracts due to increasing number of myosin bridges connecting nearby filaments [**12**]. Filaments puling the membrane cause small indentations [**13**]. These are the places where the stretching opening mechanosensitive channels is expected. In [**14**] it was shown that indeed, the cytoskeleton works as a force-transmitting and force-focusing molecular device to activate MS channels in cells. The question arises as to whether this alternative source of calcium (extracellular space) can explain so high wave velocities?

Let us notice that according to Eq.(2) the wave speed is proportional to the square root of the source amplitude A and square root of the diffusion coefficient D. The constant A tells us how intensive the source is, whereas according to the Theorem velocity can depend also on the value K of the second stable equilibrium. K is the maximal concentration at which the source term is still positive. It is also maximal possible concentration of calcium during the wave propagation. Typical values of calcium concentration in: cytoplasm, internal stores, and intercellular space are respectively equal to 10^2^ nM, 10^4^ nM, 10^6^ nM. Thus one can expect that the extracellular space can be much more intensive source of calcium than the cell internal stores. Also the possible maximal concentration of calcium at which the influx is still positive may be different then in CICR waves. There is still another mechanism proposed by L. Jaffe enhancing in some sense the diffusion of calcium – the force transmitted by the tangled filaments opens ion channels ahead of the calcium pulse and enlarges effective diffusion. Since the wave speed is proportional to square root of all these three quantities, 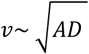 therefore, the Jaffe hypothesis may indeed be working.

## 2. Mechanically based model

The cell and tissue is often treated as the viscoelastic continuous body [**4**], which is characterized by strain and stress tensors. The stress tensor depends on the strain tensor and viscous forces dependent on the deformation speed (time derivative of strain tensor). The quantity which characterizes the mechanical action of actomyosin network is the traction force. This traction forces are resulting from the contractibility of the actomyosin network [**4**]. When calcium concentration increases the network contracts. There is some structural similarity between the thermoelastic (or thermoviscoelastic) body and the mechanical description of the cell, with traction forces of the contractile actomyosin network. Calcium similarly as heat diffuses and causes the contraction which contributes to the mechanical stress. The difference is that higher temperature leads to expansion, while higher concentration of calcium results in contraction of the cell material. Another significant difference is that the variation of calcium concentration is not resulting in the immediate appearance of traction forces. Some time (relaxation time) is needed to reorganize the actomyosin network. In general, we expect that the material of the cell represents an anisotropic elastic or viscoelastic material, similarly the traction tensor can be anisotropic. However, for simplicity we assume in this chapter both to be isotropic, so the stress tensor is characterized by two Lame coefficients and traction tensor can be expressed by a single quantity *τ*(*t*, ***x***), 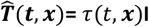 where **I** is the unit tensor, **I** = (*δ*_*ij*_).

We restrict ourselves to the linear elasticity equations. In terms of displacement field ***u***(*t*, ***x***) and stress tensor 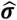 we have :

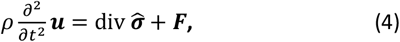

where viscoelastic stress tensor is 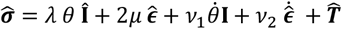 or in components

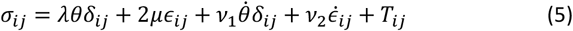

where 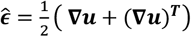 is the strain tensor, *θ* = *tr* 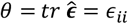 and *λ, μ* are Lame coefficients. *ν*_1_, *ν*_2_ are viscosity coefficients and *F* represents external volume forces which are vanishing in our case. Expressing Eq.(4) in terms of the displacement vector we have

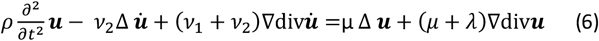

Since 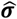 contains the traction tensor therefore we have to add the relaxation equation for the traction 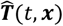 which in general anisotropic case is

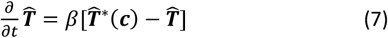

where the equilibrium traction tensor 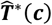 is assumed to be known function od concentration [**4**]. For the isotropic case when 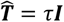, the traction is characterized by a single function *τ*(*t*, ***x***) and we have:

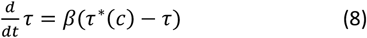

One might naively think that by solving the equations of such a model, we will determine the deformation of the membrane under the influence of the forces acting on it and, knowing the state of its stretching, we will determine the inflow of calcium into the cell. Unfortunately when the wave propagates, the cell is shrinking, its diameter gets smaller, so the continuous model do not predict the membrane stretching. Fortunately, on the microscopic level, the picture is more complex as the shrinking is caused by the actomyosin network consisting of actin filaments interlinked by myosin bridges. The number of bridges grows with calcium concentration, causing the contraction of the filament network. Fibers that are anchored to the membrane exert a force on the membrane creating funnel like wells [**13**].

**Fig. 1.**
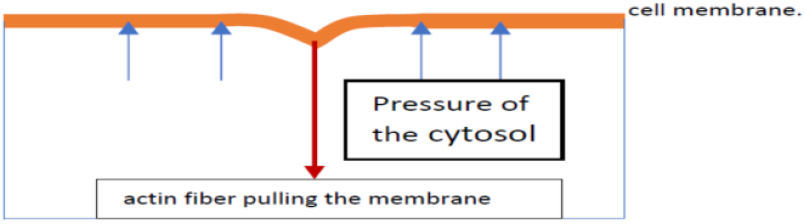
The indentations created by actin filaments.

In the areas of these indentations the membrane stretching appears. This can activate the mechanically operated ion channels. Consequently, we can have there the inflow of calcium from the extracellular space. In the language of continuum mechanics which operates with space averaged quantities the above-mentioned local stretching’s are invisible. Indeed, in the continuous limit one has the infinite number of filaments acting with infinitesimally small forces, thus the indentations of the membrane in this limit are vanishing. To estimate the possible calcium influx we have to refer to the microscopic picture of a real cell. Obviously, the number of indentations in a unit area of the membrane is proportional to the number of fibers that are anchored there, pulling the membrane. We may also assume that the part of the area where the membrane is stretched is proportional to the total force acting on the membrane. On the other hand, the average surface density of this force can be computed from the traction tensor. The important observation is that the channels cannot stay open for a long time even if stretching persists. This would lead to the enormous influx of calcium and consequently it would cause the cell apoptosis. In numerous experiments in which the membrane is locally stretched, an influx of calcium is observed (e.g. [**15,16,17**]), but the static force does not cause further influx. These remarks suggest that within a relatively short period of time the membrane relaxes and ion channels becomes inactive, which seems to agree with fluid model of the membrane. This sort of activity in the cell is observed after injury and perforation of the membrane [**18, 19,20**]. Some experiments with cells annoyed by oscillating force [**16,17**] suggest that calcium influx is proportional to the applied force magnitude and its frequency. Both arguments support the assumption that (at least in the first approximation):

### Assumption

*The calcium influx intensity is proportional to the speed of stretching not to the stretching magnitude i*.*e*., *it is proportional to the time derivative of the stretching magnitude*. Consider a unit area of the membrane with some number of indentations. Let the area of these indentation be equal to S. During stretching, the area S (t) increases. According to our considerations, the inflow of calcium through this surface should be proportional to the time derivative of S(t), i.e. to 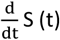. On the other hand (in first approximation) S(t) is proportional to the force applied to the membrane. In our case this force is generated by traction. In the above formulated model, the force acting on the unit membrane surface is given by *F*_*j*_ = *σ*_*ij*_ *n*_*j*_, where *n*_*j*_ are the components of the unit vector orthogonal to the surface of the membrane. If we accept the above argumentation concerning the relevance of indentations created by the actomyosin network in opening the stretched activated ion channels, then we should take into account mostly the force perpendicular to the surface of the membrane. Assuming that the cell represents the cylindrically symmetric body of radius R we have two components of this force *σ*_*rr*_ and *σ*_*rz*_. Consequently, if we assume the linear dependence then the calcium influx intensity should be expressed as

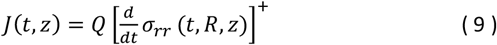

with some proportionality coefficient *Q*. Here [*f*]^+^: = max(*f*(*t, R, z*), 0) i.e. the positive part pf function *f*.

**Calcium pumps** are ion transporters located in the cell membrane [**21**]. They are responsible for active transport of calcium out of the cell, keeping the intracellular calcium concentration 10^4^ times lower than the extracellular one. There are at least two types of pumps: 1) the plasma membrane Ca^2+^ ATPase and 2) sodium-calcium calcium exchanger. These pumps and ion channels [**22**] are the main regulators of intracellular Ca^2+^ concentration. The first type is efficient at low Ca concentration, whereas the second type is extremely efficient at higher concentrations. They also play the crucial role in supporting the CICI waves. Denoting by *p*(*u*) the performance of the pumps we obtain the approximate boundary condition (used in our numerical simulations)

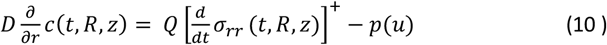

To have the equilibrium solution *c* = 0 one must assume that *p*(0) = 0, for the asymptotic stability of the ground state we will assume that *p*^′^(0) > 0. Thus (10) is a boundary condition for the calcium concentration diffusion equation

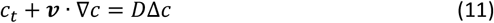

The convective term on the left (suggested by K. Piechór) represents Darcy flow of the cytosol generated by the increase of pressure during the cell compression. We have neglected here the possible contribution of calcium source from internal stores. The contribution from internal stores is rather small when compared to the boundary influx. Another reason for neglecting this term is that when performing numerical simulations, it will be known that we are dealing with pure CICI waves.

### Darcy Flow

As it is known the cytoplasm in the scale below 100nm should be considered as a liquid. The presence of sufficiently dense actomyosin filament matrix permits one for treating the cell material in larger scales as an viscoelastic gel. Therefore, in the presence of pressure gradients one can expect also the flow of cytoplasm through the porous material i.e. actomyosin matrix. Such a flow is governed by a Darcy law

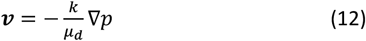

where k is the permeability and *μ*_*d*_ is dynamic viscosity od the fluid. Having cytoplasm density *ρ* as a function of pressure *ρ* = *ρ*(*p*) and the porosity *ϕ* of the filament matrix, the continuity equation (*ϕρ*)_*t*_ + *div*(***v****ϕρ*) implies

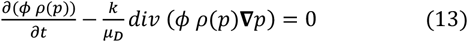

Since the variation of calcium concentration causes the reorganization of the actomyosin network, the porosity can also vary. Similarly as in the case of traction tensor we may assume that in equilibrium *ϕ* = *ϕ*^∗^(*c*) is a function of calcium concentration. If *c* = *c*(*t*, ***x***) then porosity *ϕ* is tracing its equilibrium value at each *t*, ***x*** according to the relaxation equation

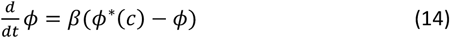

where *β* is the same as in the relaxation equation for the traction. Summarizing the mechanically based model contains following equations

1. Diffusion equation for calcium concentration with appropriate boundary condition on the lateral boundary of the cylinder (as an ideal cell).

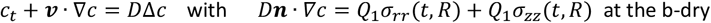
2. The mechanical equations 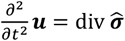 for the displacement field ***u***(*t*, ***x***) with boundary conditions relating the membrane deflection to forces acting on the membrane from the rest of the cell body, e.g. 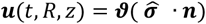, where **n** is the unit outer normal vector. To determine the deflection of the membrane is another quite complex problem of the plate loaded with some force distribution. As a very rough approach one can propose 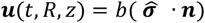 with some constant *b*.
3. The deflected membrane pushes the cytosol and generates the flow of the cytosol through the filament network which can be treated as porous material. By the Darcy law: 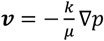 assuming that the membrane is impermeable we arrive at the boundary condition for the Darcy flow : 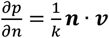, where 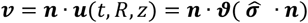, which is governed by Eqs(7-9)
4. Since 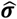 contains the traction tensor therefore we have to add the relaxation equation for the traction 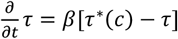 in the case of isotropic traction or Eq.(7) in general case.

Although the above system seems to fully describe the dynamics of the calcium propagation, one must mention that it still contains several simplifications, i.e. The calcium diffusion equation should take into account the existence of buffers and possible internal stores of calcium (such as endoplasmic reticula), so instead of one homogenous equation we would have the system of equations with source term representing internal stores of calcium. The influence of buffer proteins causes the substantial reduction of diffusion speed of calcium. Thus in our simplified description one has to use the “effective” diffusion coefficient which is of the order of D=30*μm*^2^/*s* whereas in water it is 10 time larger, 300*μm*^2^/*s*.

## 3. The phenomenological 3-D model

As we have seen the mechanically based model is very complex and needs a number of coefficients and material constants to be know. Therefore we still make further simplifications preserving however the main features of the model. The basic assumption is that according to Eq(10) the influx of calcium through mechanosensitive ion channels located in the membrane is proportional to the positive part of the force acting on the membrane 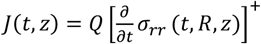. Let us estimate these forces.

### Forces acting on the membrane

To estimate the forces acting on the membrane we solve the elasticity equations making some simplifying assumptions, i.e., assuming that the membrane is stiff, remains undeformed when the actin-myosin network tries to compress the cell. The cell is represented by an infinite elastic cylinder of radius R. In addition, we will look for solution not dependent on the axial variable *z*. Neglecting the inertial term 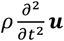, and viscous terms which as one can show are small when compared with other terms, we reduce the elasticity equations to

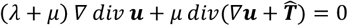

Taking into account the symmetry of the problem and 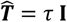, we have In cylindrical coordinates

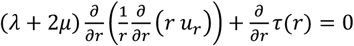

Integrating we get 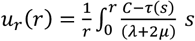 *ds*.

The boundary condition *u*_*r*_ (*R*) = 0 leads to 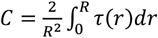. The *σ*_*rr*_ component of the stress tensor is 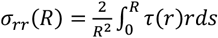 which is an average of *τ* over the cross section of the cell.

*σ*_*rr*_(*R*) is the force exerted on a unit area. Let us note that this formula was derived under assumption that *τ* is independent of *z*, therefore the fact that a force applied at some z is smeared over some neighborhood is not apparent here. one. Adopting this formula for z-dependent traction would mean that the stress generated by nonzero *τ* on some interval (*z*_1_*z*_2_) is not influencing nearby region. This is of course not true. The interconnected filaments relay the force to other region as it follows also from mechanical equations. To take it into account we will introduce some averaging operator which will be smearing out the force along *z*:

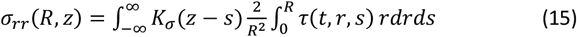

In numerical simulations we assumed *K*_*σ*_ to be the Gaussian kernel. The variance of *K* tells us what the interaction range is i.e., how far the force is relayed from one place to another. Obviously, if we knew the solution of the equation for mechanical equilibrium, we would know the range of this smearing. But it is very complex and as was already mentioned, in general, the cell can hardly be treated as an isotropic elastic material and the interactions within the filaments does not have to be correctly mapped by elastic strain and stress tensors. The range of smearing depends on degree of anisotropy. Therefore in the present work we take 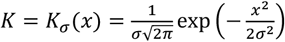 and assume that the calcium influx through the ion channels per unit area of the lateral boundary according to Eq.(9) is given by

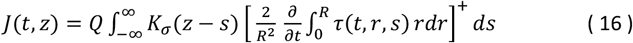

Although in deriving this formula we made several simplifications we believe that the possible discrepancy can be accommodated in the proportionality constant *Q*.

The last formula is not yet the full boundary condition for calcium diffusion equation. There is still very important mechanism that must be taken into account. Once the calcium enters the cell it must be somehow evacuated soon to restore the equilibrium. As was already mentioned this role is played by special pumps located in the membrane. They are pumping out the calcium, thus restoring the cell equilibrium. As the boundary condition for diffusion equation, we take the expression

Since the integral with respect to *s* in Eq(16) is the convolution with respect to z variable and the time derivative of *τ* can be expressed by the right hand side of Eq.(16) therefore we can write

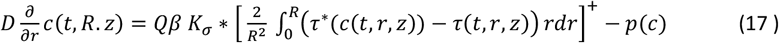

where *p*(*c*) is the pumping intensity.

In the calcium diffusion equation we will neglect the convective term, which is expected to be rather small in comparison with other terms so we have

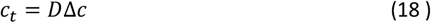

To close the model, we have to add the relaxation equation for the traction

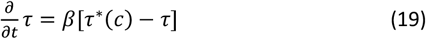

Thus our simplified model consists of two equations, Eqs(18,19) and the boundary condition Eq(17) The constant *Q* tunes the intensity of calcium inflow. However, as will follow from the analysis of 1D case, the traveling wave solution exists if the “reduced” source term *τ*^∗^(*c*) − *p*(*c*) is a bistable function. Therefore, it is convenient to introduce just one constant defining the intensity of the whole source term as was the constant A in example (2). As far as *p*(*c*) is not defined it does not make any difference, but now varying A we will preserve that *A*(*τ*^∗^(*c*) − *p*(*c*)) is of a bistable type. For this reason, we write the influx boundary condition in the form

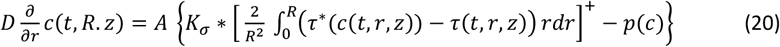

Let us comment the importance of the + sign in the above boundary condition. As follows from numerical simulations, once it is omitted, the solution for calcium concentration takes also negative values.

The system (18-20) was studied numerically. The travelling wave solutions vanishing in both infinities were found if β was sufficiently small. When β was growing the with of the wave was decreasing as 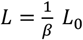 When the rear of the wave was approaching its front and *L* becomes comparable to the sum of the front and rear thickness the wave disappeared.

## 4. One dimensional approximation

Now, assuming that the radius R is not too large we will derive a averaged one dimensional approximation of our problem. Let us denote 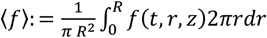 for any function *f* and let

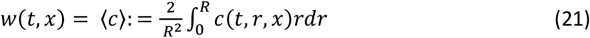

To derive the 1-D approximation we make the averaging of our diffusion equation with respect to r :

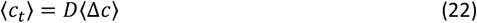

We have ⟨*c*_*t*_⟩ = *w*_*t*_ and 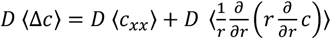. The last term on the right can be expressed in terms of the influx of calcium through the boundary Eq.(20)

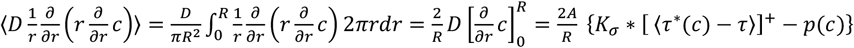

For *τ*^∗^(*c*) = *a c*, i.e. depending linearly on *c* we obviously have ⟨*τ*^∗^(*c*)⟩ = *τ*^∗^(*w*). In case of nonlinear dependence this can be assumed as a reasonable approximation for not too large R. In case of the expression *p*(*c*) = *p*(*c*(*t, R, z*)), which represents the action of pumps located in the membrane the concentration *c* should be taken at the boundary, so again *p*(*c*(*t, R, z*)) ≈ *p*(*w*) is an approximation. With the above approximations we arrive at one dimensional problem

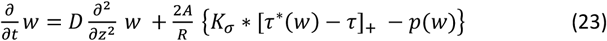

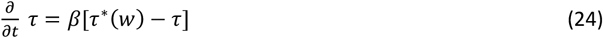

where 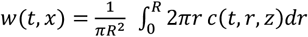In the limiting case of short range interaction 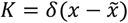 one obtains

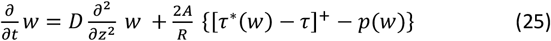

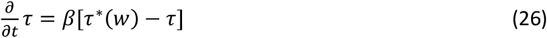

which is of a FitzHugh -Nagumo type [**23**] with somewhat untypical source function. As will be seen later we will derive another still simpler 1-D approximation similar to Eqs(25-26) but with modified diffusion.

## 5. Numerical simulations

The 3-D system (-) was solved numerically assuming that *τ*^∗^(*c*) is equal to

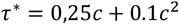

and *K*_*σ*_(*z*) is of gaussian type. For *p*(*c*) which models the action of pumps we took as an example:

*p*(*c*) = *c*(*c*^2^ − 1.15*c* + 0.5). As we said before, the existence of two types of pumps is essential for the existence of calcium waves. The first ones are efficient at low concentrations - the pumping rate increases with increasing concentration and then stabilizes, and the other is very efficient at high concentrations. Therefore, we may expect that for some *C* = *c*_*in*_ there is an inflection point of p(c). Thanks to this, there is a possibility that *f*: = *τ*^∗^(*c*) − *p*(*c*) becomes a bistable function. The source term for τ=0 and *K*_*σ*_ = *δ*(*x*) (see Eq.(25)) up to multiplicative constant

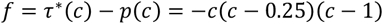

is bistable function. Consequently, in 3D simulations as the boundary condition for the calcium diffusion equation we have for r=R:

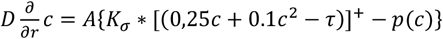

Where 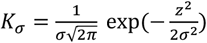. All numerical computations were done for the diffusion coefficient D=1 and 1) A=1 in 1D cases, whereas 2) in 3D simulations we used A=1 for all cases with R=2, and A=20 for cases with R=10,20,40. Let us notice that for σ=0, β=0, hence τ=0, Eqs(23,24) are reduced to the reaction diffusion equation 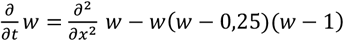 which has heteroclinic solutions (travelling fronts) of the form 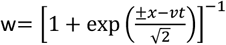 moving with velocity 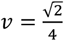. All velocities on the Fig.1 are related to this velocity. We expect that for *σ*=0 the velocity of travelling wave for our problem should be close to this velocity. The thickness of the front is small when compared to the wave width and τ at the wave front is closed to zero so the heteroclinic solution is there a reasonable approximation.

**Fig. 2.**
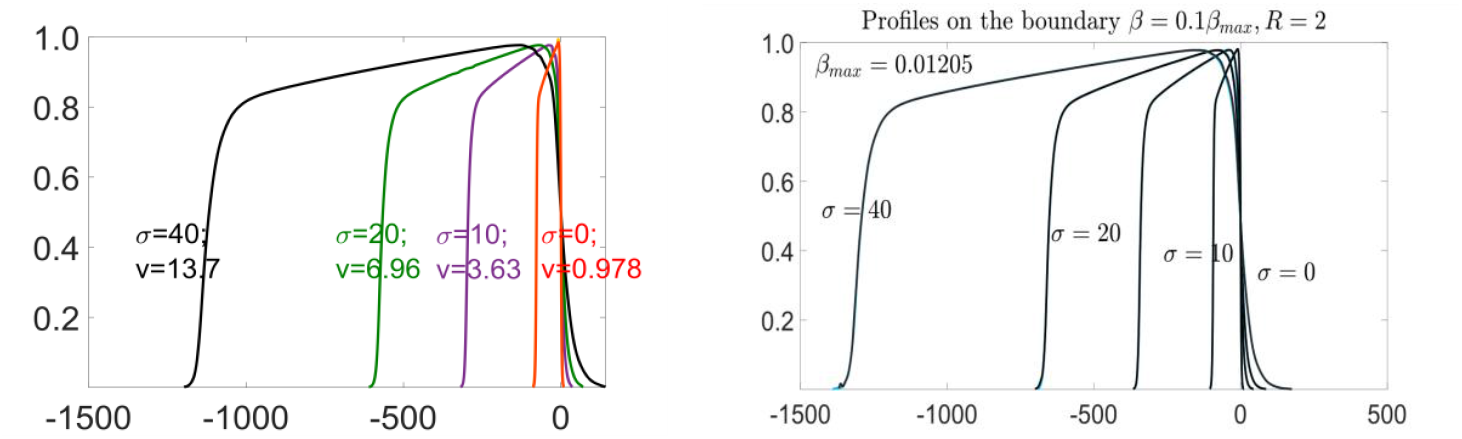
On the left: wave profiles and wave velocities in 1-D simulations for A=1, and *σ*= (0,10,20,40) and for *β*=0.001515. On the left: 3D simulations for A=1, R=2, and slightly smaller β, β=0,001205 and the same values of σ.

As expected for σ=0, the wave velocity is almost the same as in the case *β*=0 and it grows with σ. Similarly, the propagation velocity also grows with σ. As can be seen from Fig.2, there is very good agreement between the 1-D and 3-D simulations. The wave widths are slightly larger on the right picture, mainly because of the difference in β.

Below some results of numerical simulations in 3-D for A=20 and starting from the top for β= 0.01205, β = 0.0241, β= 0.03615 are presented. The wave is moving from the left to the right. Similarly as in 1-D case the width of the pulse seems to behave as 1/β. Please note that for β=0 the pulse becomes infinitely long of heteroclinic type.

In the last figure the wave profiles are presented for A=1, R=2, σ=20 and different β. One can notice that the width of the wave is inverse proportional to β. 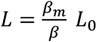, where for *L*_0_ we can take approximately 2×(thickness of the front).

**Fig. 3.**
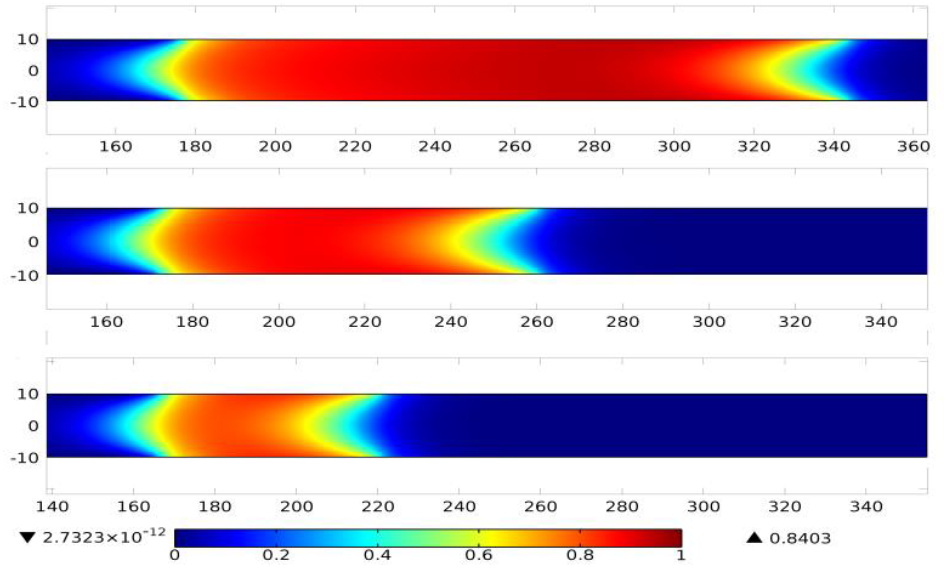
Examples of numerical solutions of the waves running from the left to right for *R* = 10, σ=10 and different *β*; starting from the top: β=0.1*β*_*max*_, β=0.2 *β*_*max*_, β=0.3 *β*_*max*_, where *β*_*max*_ = 0.01205

**Fig. 4.**
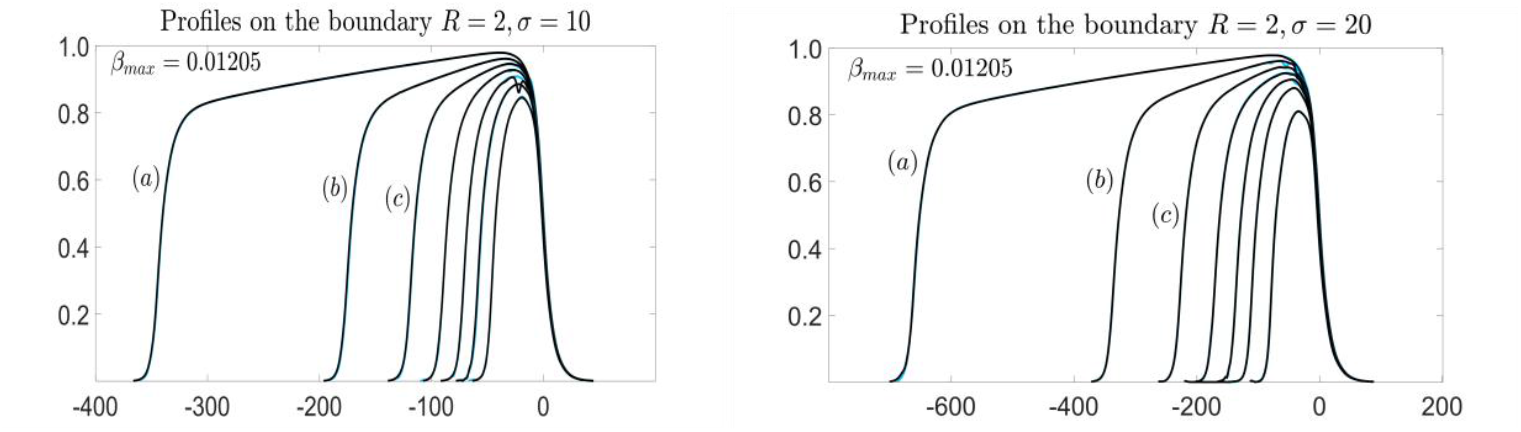
Waves profiles at r=R, (R=2) for different β: a) β=0,1*β*_*max*_, b) β=0,2*β*_*max*_ c) β=0,3*β*_*max*_ etc. where *β*_*max*_ = 0,01205. On the left for σ=10. On the right for σ=20.

## 6. The influence of the variance *σ* of *K*_*σ*_ on the wave velocity

For *β* = 0 the travelling wave solution of 1D approximation has the form of heteroclinic front which connects stable equilibria *w*_1_ = 0 and *w*_3_ = 1. For very small *β* the front part of the solution is almost identical with that for *β* = 0, but later the value of the pulse falls down, at first slowly and then much faster, from *w* = 1 to w =0 at infinity. Therefore, the wave velocity is practically the same as it is for the heteroclinic case, *β* = 0. In this case however, we have *τ*^∗^(*w*) ≥ 0 so [*τ*^∗^(*w*)]_+_ = *τ*^∗^(*w*) and the system reduces to

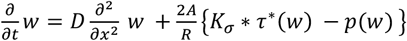

We have

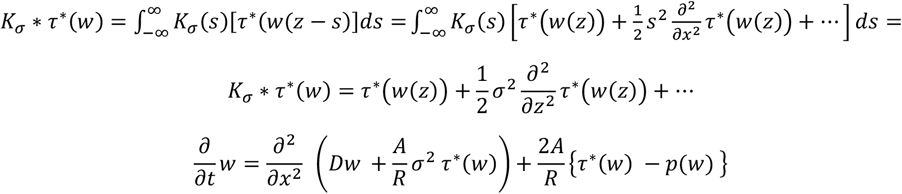

Thus, for small *β* the front part of the wave can be described by the reaction diffusion equation with the bistable function. If *τ*^∗^(*w*) ≅ *a w* as in (), then the propagation velocity is given by (see Eq.(3) where D has to be replaced by 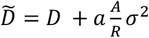, hence 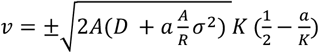. For values that were taken in numerical simulation i.e. *D* = 1, *a* = 0.25, *K* = 1 we have

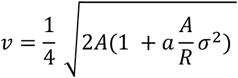

This formula seems to be in a good agreement with numerical simulations for sufficiently small β. When β grows the wave slows down as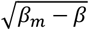 where *β*_*m*_ is the maximal value of β for which the traveling wave solution exists. *β*_*m*_ depends on σ, D and A.

## 7. The FitzHugh-Nagumo type of approximation

Extending the above approximation also to this part of the wave where τ is not small and cannot be neglected we come to the following system of equations

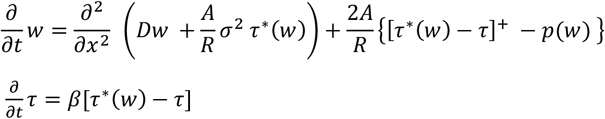

This system (especially for linear *τ*^∗^(*w*)) is very close to FitzHugh-Nagumo equation. The essential difference is that here we take [*τ*^∗^(*w*) − *τ*]^+^ - the positive part of (*τ*^∗^(*w*) − *τ*) which guarantees positivity of *w*(*t, z*) for positive initial values of *w*. This system is much simpler than the nonlocal one i.e. Eq(23) and Eq(24) of 1-D approximation although it provide us with quite similar results.

## Discussion

In [13] point like forces of 5 - 20 of pN (piconewtons) were applied to deform the cell membrane. The forces in this range are comparable with the forces exerted by a single myosin head (in muscle motor myosine), which are about 3-6 pN (see e.g.[24] p.381) or the forces generated by growing microtubules [25]. The depth of the corresponding indentation evoked by such forces is of the order of 500 nm. The experiments used mouse neuroblastoma rat glioma hybrid (NG108-15) line of cells. The results of the experiments performed in [13] justify, at least partially, the assumptions of our model. The contracting network of actin filaments puling the membrane causes the number of indentations, thus stretching the membrane. This opens the mechanosensitive ion channels. Assuming that the intensity of the inflow of calcium is proportional to the positive part of the time derivative of the force exerted on the membrane we derived the phenomenological mathematical model of calcium waves. Their propagation is supported by the influx of calcium to the cell from extracellular space. The basic model is an analogue of a 3-D nonlocal FitzHug-Nagumo system. Averaging this system over cross-section of the (ideal cylindrically symmetric) cell we derived nonlocal 1-D approximation.

It turns out that the 1-D system can be still approximated by a local equation – a sort of the Fitzhugh-Nagumo with a specific source term. Numerical simulations show that solutions of all these three systems have nonnegative traveling wave solutions – travelling pulses with a single extremum. Although the discrepancy between solutions of 1-D and 3-D (in the width or velocity of the wave) grows with R, the agreement seems to be quite good.

## Acknowledgments

The research was supported by: National Science Centre (Poland) grant 2016/21/B/ST1/03071.

Supplements. One should study the full symmetry for the system.

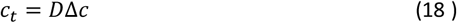

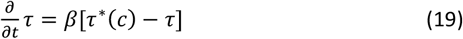

with the boundary condition on r=R (where we suppose the membrane is located)

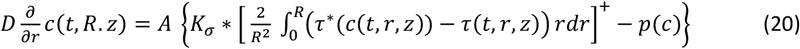

Where 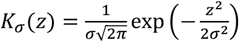.

2. Symmetries for 1D model

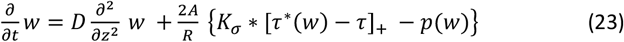

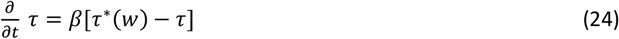

where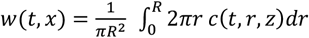

In the limiting case of short range interaction 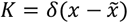 one obtains

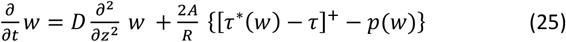

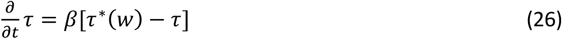

We have

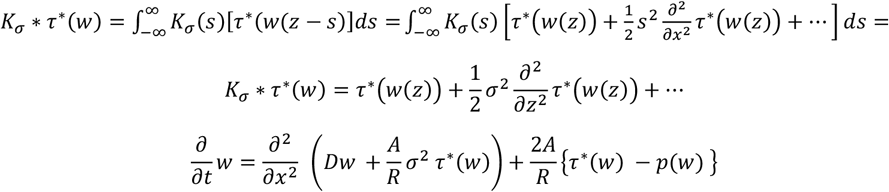

Thus, for small *β* the front part of the wave can be described by the reaction diffusion equation with the bistable function. If *τ*^∗^(*w*) ≅ *a w* as in (), then the propagation velocity is given by (see Eq.(3) where D has to be replaced by 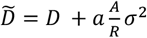

